# Population suppression by release of insects carrying a dominant sterile homing gene drive targeting *doublesex* in *Drosophila*

**DOI:** 10.1101/2023.07.17.549342

**Authors:** Weizhe Chen, Jialiang Guo, Yiran Liu, Jackson Champer

## Abstract

Gene drive alleles, which bias their own inheritance and increase in frequency, show great promise for blocking disease transmission or directly suppressing pest populations. The most common engineered drive system is the CRISPR homing drive, which converts wild-type alleles to drive alleles in the germline of drive heterozygotes by homology-directed repair after CRISPR cleavage. One successful homing drive example targets a female-specific exon in *doublesex* in *Anopheles* mosquitos, suppressing the population by inducing recessive sterility in female drive homozygotes. We found that in *Drosophila melanogaster*, a 3-gRNA drive disrupting the *doublesex* female exon resulted in a masculine phenotype and dominant female sterility. Resistance alleles formed by end-joining repair were also dominant sterile. This was likely caused by expression of male-specific transcripts in females with drive and resistance alleles, disrupting sex development. Based on this construct, we proposed a new pest suppression system called “Release of Insects carrying a Dominant-sterile Drive” (RIDD). This entails continuously releasing drive heterozygous males, with drive and resistance alleles causing sterility in females. The drive remains at high frequency longer than currently used dominant female-lethal alleles (RIDL) due to drive conversion in males, and drive alleles also cause sterility based on resistance, both substantial advantages. With weekly releases of drive males into a cage population with overlapping generations, our RIDD system targeting *dsx* reached 100% prevalence within 27 weeks, progressively reducing egg production and eventually causing total population collapse. RIDD combines the merits of homing gene drive and RIDL. It is powerful but self-limiting, unlike unconfined standard homing drives, allowing for targeted population suppression.

## Introduction

Insect pests cause vast damage to agriculture and pose threat to public health ^1, 2^. Chemical pesticides have been utilized to suppress pests for decades, but increasing development of pesticide-resistance and harmful environment effects prevent this from being sustainable ^3–5^. Thus, effective and ecologically friendly pest control techniques are urgently required. In response to this growing need, a number of pest control technologies have developed in recent years based on genetic engineering biotechnology ^6–11^. Some of these strategies have successfully suppressed target populations in laboratory trials, which can potentially lead to deployment in the future ^7, 12–15^.

Sterile insect technique (SIT), an environmental-friendly and species-specific pest control method, was the first genetic pest control system ^16^. It aims to reduce local reproductive potential and thereby reduce the population. A large number of male insects sterilized by radiation or other methods are released to the wild, mating with wild females, which then produce no or fewer offspring ^6, 10, 16^. If sufficient sterile insects are released repeatedly, the local population will be reduced or even eliminated. SIT programs have been conducted with varying degrees of success against screwworm, tsetse, medfly, and *Cochliomyia hominivorax* ^6, 10, 17, 18^. However, despite its attractive features, exposing insects to sterilizing doses of radiation or chemicals reduces their longevity and mating competitiveness ^19^. This means that in many scenarios, it may be impractical to release enough insects to cause population suppression.

With the development of genetic engineering, transgenic pest control strategies were proposed to improve and complement conventional SIT. Female-specific release of insects carrying a dominant lethal gene (fsRIDL) system is similar to SIT, avoiding the side effects of irradiated-sterilization by using genetic manipulation ^18–21^. Dominant lethal transgenes specifically expressed in females yield several advantages compared with SIT. The transgenic allele can be transmitted by released transgenic males, rendering the female offspring nonviable. The heterozygous male offspring carrying the transgenic allele can continue to help suppress the population by transmitting it to half of their daughters. Thus, not only do fsRIDL males potentially have a fitness advantage over SIT males (due to lack of harmful sterilization technique), they also can cause effects over multiple generations, making the technique substantially more powerful than SIT^22, 23^. It avoids the need for sex separation and radiation/chemosterilization as well, though rearing may still be more expensive than SIT due to the need to use a ligand to repress the female-lethal allele. fsRIDL has been utilized to successfully suppress fruit flies ^24^, mosquitos ^19, 25–28^, and moths ^29, 30^ in cage or field trials. Nevertheless, fsRIDL still relies on mass-rearing technology, which is costly and demanding in some species ^14, 25, 31–35^, so it may still often lack the power necessary for success.

At the frontier of pest control, gene drive shows great promise for blocking disease transmission or directly suppressing pest populations ^7, 8, 12, 15, 36^. Gene drives are alleles that can bias their own inheritance, increasing in frequency and eventually spreading through a population along ^37^. The most common engineered gene drive system is CRISPR homing drive, which converts wild-type alleles to drive alleles in the germline of drive heterozygotes. Specifically, the target site is cleaved by Cas9 and a guide RNA (gRNA), and the DNA break then undergoes homology directed repair (HDR). To accomplish population suppression, CRISPR homing drive can be designed to target a haplosufficient but essential gene, still allowing drive conversion to take place in healthy heterozygotes, while reducing the fertility or viability of homozygous individuals ^38^.

One potential target is *doublesex* (*dsx*), which is a central nexus for sex determination in insects ^22, 39, 40^. With alternative splicing, sex-specific *dsx* transcripts are necessary for sexual differentiation^41^. Females that are homozygous for a disrupted *dsx* allele at the female-specific exon (by insertion of drive allele or a nonfunctional resistance allele) lose reproductive capability. Kyrou et.al successfully suppressed a cage population of *Anopheles gambiae* by targeting the intron 4/exon 5 junction of *dsx* ^40^.

High efficiency homing drive enables population elimination with a small initial release if the drive is sufficiently efficient. However, this is a double-edged sword: the zero-introduction threshold implies that the drive cannot be confined to a target population because even small numbers of migrants will spread the drive to other even weakly connected populations, which may raise sociopolitical concerns or be undesired from an ecological point of view ^9, 36, 42^.

We here propose a strategy, Release of Insects carrying a Dominant sterile Drive (RIDD), combining advantages of homing gene drive and RIDL. We found that disruption of the *dsx* female-specific exon is dominant sterile in *Drosophila melanogaster* females, either by the drive or by most nonfunctional resistance alleles. The drive has no apparent impact on males, and drive conversion can take place in germline of male heterozygotes. Thus, when drive males are released to wild population, the drive allele can spread when they mate with wild-type females. The male offspring that carry a drive allele can continue gene drive in the next generation, while female offspring with a drive allele or resistance allele, which together compose all transmitted alleles, will be sterile. Repeatedly releasing RIDD males into cage populations resulted in slow increase in drive allele frequency and eventually population elimination after several generations. Modeling indicates that RIDD can be substantially more efficient than RIDL, suggesting its potential use in future pest control.

## Methods

### Plasmid construction

Our plasmids construction was based on TTTgRNAtRNAi, TTTgRNAt, and HSDygU4, which were constructed previously. *dsx*-related fragments were amplified from the genome of *w*^1118^ flies. We selected gRNA target sites from the website CHOPCHOP. Reagents for restriction digest, PCR, and Gibson assembly, plasmid miniprep were obtained from New England Biolabs and Vazyme; primers were from Integrated DNA Technologies Company BGI; 5-α competent *Escherichia coli* from Vazyme; and the ZymoPure Midiprep kit from Zymo Research. Plasmid construction was confirmed by Sanger sequencing. Detailed information about DNA fragments, plasmids, primers, and restriction enzymes used for cloning of each construct are listed in the Supplementary material. Plasmid sequences of the final drive insertion plasmid and target gene genomic region with annotation are provided on GitHub (https://github.com/jchamper/ChamperLab/tree/main/dsx-Suppression-Drive) in ApE format ^43^.

### Generation of transgenic lines

Embryo injections were conducted by Fungene Transgenic Flies Company. The donor plasmid HSDdsxRed3g (300ng/ul) was injected into *w*^1118^ flies together with TTChsp70c9 (300ng/ul) providing Cas9 for transformation. Flies were housed in modified Cornell standard cornmeal medium (using 10 g agar instead of 8 g per liter, addition of 5 g soy flour, and without the phosphoric acid) in a 25℃ incubator on a 14/10-h day/night cycle at 60% humidity.

### Phenotypes and Morphological Analysis

Flies were anesthetized with CO_2_ and screened for fluorescence using the NIGHTSEA adapter SFA-GR for DsRed and SFA-RB-GO for EGFP. Fluorescent proteins were driven by the 3×P3 promoter for expression and easy visualization in the white eyes of *w*^1118^ flies. DsRed was used as a marker to indicate the presence of the split drive allele, and EGFP was used to indicate the presence of the supporting *nanos*-Cas9 allele. Morphological photos were taken by phone using stereo microscope with 10×/22 magnification.

### Male mating competitiveness assay

For preparing drive and non-drive males with similar fitness, their mothers were *nanos-*Cas9 sisters of similar age, and they were allowed to lay eggs in the same vial. In each vial, we added one Cas9 female that was previously mated with a Cas9 male, and another that was mated with a drive/Cas9 male (all these flies were homozygous for Cas9). These were allowed to lay eggs for three days and were then removed. After the offspring hatched, we phenotyped and paired one drive/Cas9 male with one Cas9 male of the same age for a competition test. For each test, we added one *w*^1118^ female, one drive/Cas9 male, and one paired Cas9 male to a vial and allowed mating for one day. Then, we removed the males and allowed the female to lay eggs for several days. We phenotyped the offspring, and if the percentage of individuals with red fluorescent eyes was higher than approximately 60-70%, we scored this as a drive male mating success. If all the offspring were wild type, we scored this situation as a *w*^1118^ mating success. Intermediate levels of drive inheritance between these indicated that both males likely mated with the female and were scored as a tie.

### Cage study

For the cage study, flies were housed in 25×25×25cm enclosures with 7 food bottles at first with the oldest bottle being replaced every two days (a fourteen-day cycle), and after four weeks of drive releases was gradually (over two weeks) reduced to 5 bottles with one bottle being replaced every three days (a fifteen-day cycle). Flies that were homozygous for the split *nanos*-Cas9 allele were initially added to each cage, with an initial group of several hundred of varying ages. They mated freely and laid eggs in food bottles for eight weeks in the cage to establish to initial cage population, which was observed to reach an equilibrium density after approximately 4-5 weeks. A control cage was kept on the 15-day cycle without any addition of drive flies. For the drive test cages, drive males were released every week, and the first week was counted as “week zero”. The drive line was heterozygous for the split *dsx* homing drive allele and homozygous for the *nanos*-Cas9 allele, and it was generated by crossing drive males to females with the *nanos*-Cas9 line for several generations, selecting males with brighter green fluorescence (which were more likely to be Cas9 homozygotes). To continuously generate drive male flies for release, we set up bottles with crosses between drive/Cas9 males and Cas9 females. Drive male flies were collected and released into the cage each week (usually in two evenly spaced batches) after the initial cage population was established. Detailed release numbers of each week is shown in the Supplementary Data.

To record the dynamics of the cage population, we took photos each week of each side of cage to estimate the total population, and we randomly aspirated some flies from the cages for phenotyping. These flies were then returned to the cage. Also, to estimate the allele frequency of new hatching flies in the cage, we kept some removed cage bottles for one day and then phenotyped any new adults that emerged during this day.

### Phenotype data analysis

Pooled analysis was conducted for calculating drive inheritance, drive conversion and other parameters by combining all data from different individual crosses. However, this pooling approach does not consider potential batch effects (each individual cross is considered as a separate batch with different parameters, which could bias rate and error estimates). To account for such batch effects, we conducted an alternate analysis as in previous studies^44, 45^. In brief, fitting a generalized linear mixed-effects model with a binomial distribution (maximum likelihood, Adaptive Gauss-Hermite Quadrature, nAGQ = 25) enables variance between batches, which then results in marginally different parameter estimates but higher standard error estimates. This analysis was performed with R (3.6.1) and supported by packages lme4 (1.1-21, https://cran.r-project.org/web/packages/lme4/index.html) and emmeans (1.4.2, https://cran.r-project.org/web/packages/emmeans/index.html). The code is available on Github (https://github.com/jchamper/ChamperLab/tree/main/dsx-Suppression-Drive). In our study, the alternate rate estimates and errors were close to the pooled analysis (Supplemental Data).

### Diagnostic PCR

To figure out transcription of *dsx* in drive and resistance allele carriers, flies were frozen and homogenized. RNA was extracted by using RNeasy Mini Kit, and reverse transcription was used to obtain cDNA with RevertAid First Strand cDNA Synthesis Kit with oligo(dT) primers. This cDNA was the template for PCR using Q5 DNA Polymerase from New England Biolabs with the manufacturer’s protocol. Primers Exon3_S_F and Exon4_S_R were designed to specifically amplify the female-specific *dsx* transcript, and Exon3_S_F and Exon5_S_R were designed to amplify the male-specific *dsx* transcript.

### Sanger Sequencing

Flies were frozen, and DNA was extracted by grinding single flies in Trizol DNA extraction Reagent. The DNA was used as a template for PCR using Q5 DNA Polymerase from New England Biolabs with the manufacturer’s protocol. To sequence resistance alleles, the region containing the gRNA target sites was amplified using DNA oligo primers dsRed_S_F and dsx_S_R. To determine the sequence of drive allele elements in individuals with faint red fluorescent eyes, the DsRed region was amplified by the primer pairs dsx_S_F/CFD5_S_R and dsx_S_F/dsRed_S_R. After DNA fragments were isolated by gel electrophoresis, sequences were obtained by Sanger sequencing and analyzed with ApE software.

### Population models

Deterministic, discrete-generation models assume that only wild-type females would reproduce (any with drive or dominant-sterile resistance alleles are unable to) by randomly selecting a mate from the pool of available native males or continuously released males. Females then generate offspring based on the male genotype. In drive heterozygous males, wild-type alleles are converted to a drive allele in the germline at a variable efficiency, and remaining wild-type alleles are assumed to be converted to dominant female-sterile alleles. At the end of a generation cycle, offspring genotype frequencies are normalized to generate the final allele frequencies. This model was built in Excel and is available on GitHub (https://github.com/jchamper/ChamperLab/tree/main/dsx-Suppression-Drive).

Stochastic simulations were performed in SLiM 4.0. ^46^ similar to previous studies^47–50^. Our simulations have a single panmictic population of diploids with discrete generations. Each individual is specified by its genotype, which can have wild-type alleles, drive alleles, and resistance alleles. Drive alleles and resistance alleles cause dominant sterility in females, but we assume no other fitness costs. Drive/wild-type heterozygous males will convert a fraction of wild-type alleles in their germline into drive alleles at the drive conversion rate. Remaining wild-type alleles are all converted to resistance alleles.

In any generation, the population is defined by the numbers of male and female adults of each genotype. To determine the genotypes in the next generation, each female first selects a random male in the population to mate with. This could be a native-born male, or a newly released male, with drive heterozygous males being released at a fixed number in each generation. After a mate has been selected, the female will have a number of offspring drawn from a distribution with an average of 2 * [(1-β) * N / K + β) where β is the low-density growth rate, N is the current population size, and K is the carrying capacity of the population. As default values, we used β = 10 and K = 50,000. The actual number of offspring for an individual female is determined by a draw from a binomial distribution with 50 trials, yielding a maximum of 50 offspring and an average of 2 offspring for a wild-type female when the population is at its carrying capacity. Each offspring generated is assigned a random sex, and its genotype is determined by randomly selecting one allele from each parent, with adjustments for drive activity. SLiM files used in this study are available on GitHub (https://github.com/jchamper/ChamperLab/tree/main/dsx-Suppression-Drive).

A SLiM framework was also used to simulate the dynamics of RIDD in our *Drosophila* cage populations. In this model, time steps were weekly. Flies suffered density-dependent mortality at the egg stage at age 0 as (2 * β / 100) * X / [(β –1) * N / K + 1), where N is the population size, and K is the expected carrying capacity. X is a constant that produces an appropriate survival level to maintain the population at carrying capacity, determined by the age-based death rate. The population capacity was set to 2500. Females produced an average of 100 eggs per week, drawn from a Poisson distribution. Larvae that were 1 week old suffered no mortality (mortality for these was factored into the mortality for age 0 larvae in the model). Adult mortality was set so that an age class linearly declined to zero by week 9 for males, and female survival was set as 80% of male survival to account for higher mortality during egg laying (where overcrowding was observed to produce high death rates, resulting in a male-skewed sex ratio). 100 drive males were released into a panmictic population each week for first ten weeks, after that the release size increased to 600 per week. The low-density growth rate was 10, roughly estimated based on average egg laying rates, survival, and time for hatching before removal of food. Because the effective population size in our experimental cages was likely lower than the censuses sizes, the actual simulation results we show have all population sizes (listed above) divided by ten noted above to produce stochastic variation based on typical effective population sizes^51^.

## Results

### 1. Homing drive design targeting *dsx*

We aimed to develop homing suppression drive in *D. melanogaster* by targeting *doublesex*. Null mutations of *dsx* results in female recessive sterility but show no effects on males, according to the previous drive study in *Anopheles* mosquitoes ^40^. *dsx* is a crucial central “nexus”, responding to upstream sex determination signals as well as regulating thousands of downstream sexual differentiation cascades ^22, 39^. A highly conserved female-specific alternative splicing acceptor site is near the boundary of intron 3 and exon 4, resulting in different *dsx* transcripts in males and females ^52, 53^ (Fig.2c).

Our drive is inserted between the leftmost and rightmost gRNA target sites of *dsx* (Fig. 1a). DsRed driven by 3×P3 promoter is included as a fluorescent marker to indicate the presence of a drive allele. The drive also contains three gRNAs, all expressed by the U6:3 promoter and multiplexed by tRNA sequences, avoiding repetitive gRNA promoter elements. The gRNAs target near the boundary between intron 3 and exon 4, with all cut sites within the exon (though the first cut site is only two nucleotides into the exon). These targets ensure that frameshift mutations or disruption of sexual alternative splicing of the female-specific exon would firmly disrupt the gene’s function in females. Our gRNA sites were also chosen to avoid strong off-target sites.

**Figure 1.**
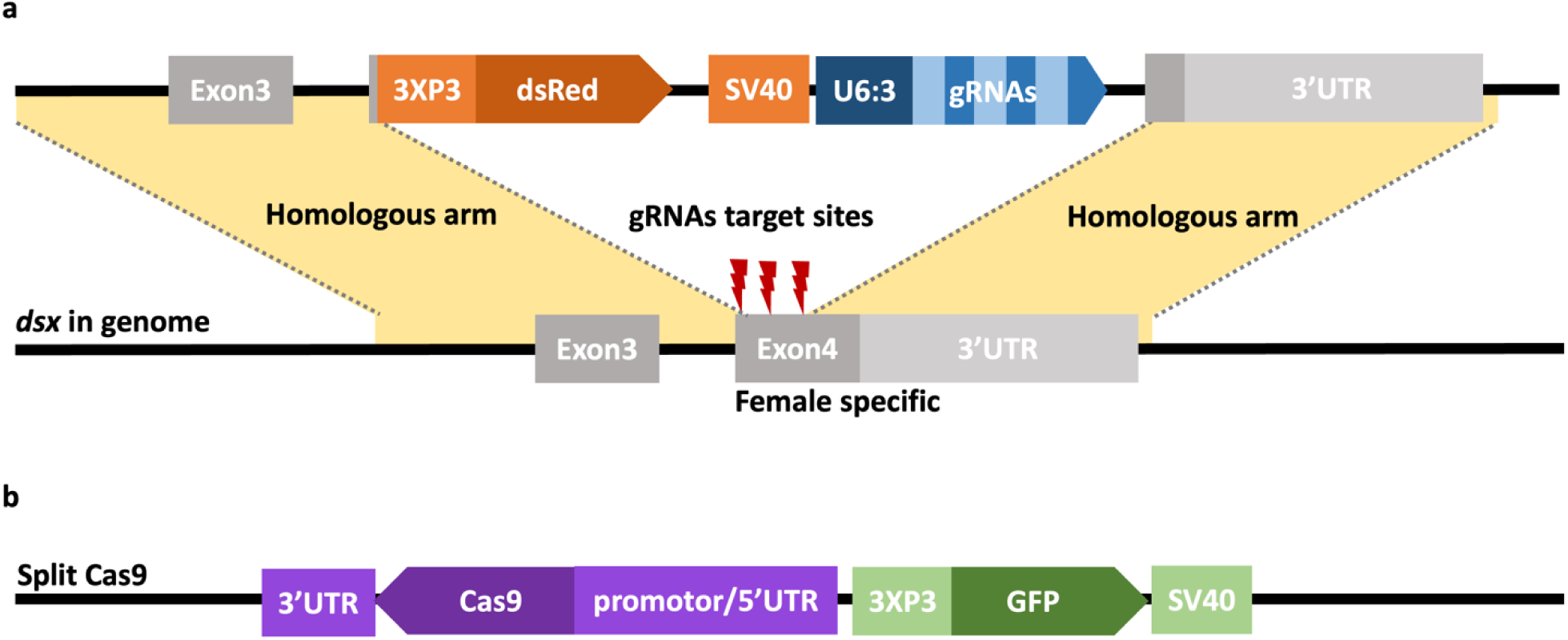
Gene drive constructs. (**a**) Our gene drive construct consists of a dsRed fluorescent marker driven by the 3×P3 promoter and three gRNAs driven by the U6:3 promoter that are separated by tRNAs. Exon-4 of *dsx* on chromosome 3R is female-specific exon. gRNAs target the intron 3/exon 4 boundary within the exon. In a drive heterozygote’s germline, Cas9 and gRNA cleaves the wild-type allele. Repair of the cleaved chromosome through homologous directed repair (HDR) leads to copying of the drive allele (called “drive conversion” or “homing”). (**b**) The split Cas9 construct is on chromosome 2R. Cas9 is under the control of is one of the following 5’UTR/promotors: *nanos*, *rcd-1r* and CG4415. The 3’ UTR is either *nanos* or *shu*. A 3×P3:EGFP cassette is used to indicate the presence of the Cas9 allele.

The Cas9 element, required for drive activity, is placed on chromosome 2R and provided through a separate line carrying Cas9 and EGFP with the 3×P3 promoter. To assess the effect of Cas9 regulation on drive performance, we tested our drive with several Cas9 constructs (Fig. 1b), which is comprised of different 5’UTR and 3’UTR. The main Cas9 line used *nanos* regulatory elements ^54^, but we also tested combinations of CG4415 and *rcd-1r* promotor/5’UTR elements with *nanos* and *shu* 3’UTR elements, which were recently shown to support high drive performance^55^. The drive will only be active in individuals both containing Cas9 and drive allele in this split drive system ^54^.

### 2. Drive performance and morphological analysis of the *dsx* drive

To assess the drive conversion efficiency, we crossed *dsx*-drive male heterozygotes with five different Cas9 lines females, differing in Cas9 regulatory elements. The offspring with both green and red fluorescent eyes, indicating that they were heterozygous for the drive allele and Cas9 allele, were used for drive performance and morphology assessment.

For males, the *dsx* drive shows no impact on their morphology and fertility. Drive conversion (also called homing) takes place in the germline of heterozygotes, where the wild-type allele can be converted to a drive allele by CRISPR cleavage and homology-directed repair (Fig. 2a). In crosses between drive and Cas9 heterozygous males and *w*^1118^ females, the drive allele was inherited at a high rate, up to 85% with Cas9 driven by the *nanos* promotor or *rcd-1r* promotor (84.8% with *nanos* promotor/*nanos* 3’UTR, 85.5% with *rcd-1r* promotor/*nanos* 3’UTR, 81.8% with *rcd-1r* promotor/*shu* 3’UTR). Drive inheritance was bit lower with the *CG4415* Cas9 promoter, but still at least 70% (70.0% with *CG4415* promotor/*shu* 3’UTR and 73.3% with CG4415 promotor/*nanos* 3’UTR) (Fig.2a). These drive inheritance rates were all significantly higher than the Mendelian inheritance rate of 50% (Fisher’s exact test, *P* < 0.0001) and thus indicative of strong drive activity in the male germline. For the *nanos* promoter/5’UTR/3’UTR combination, the drive conversion rate was 69.6%, assuming that all genotypes had the same viability.

**Figure 2.**
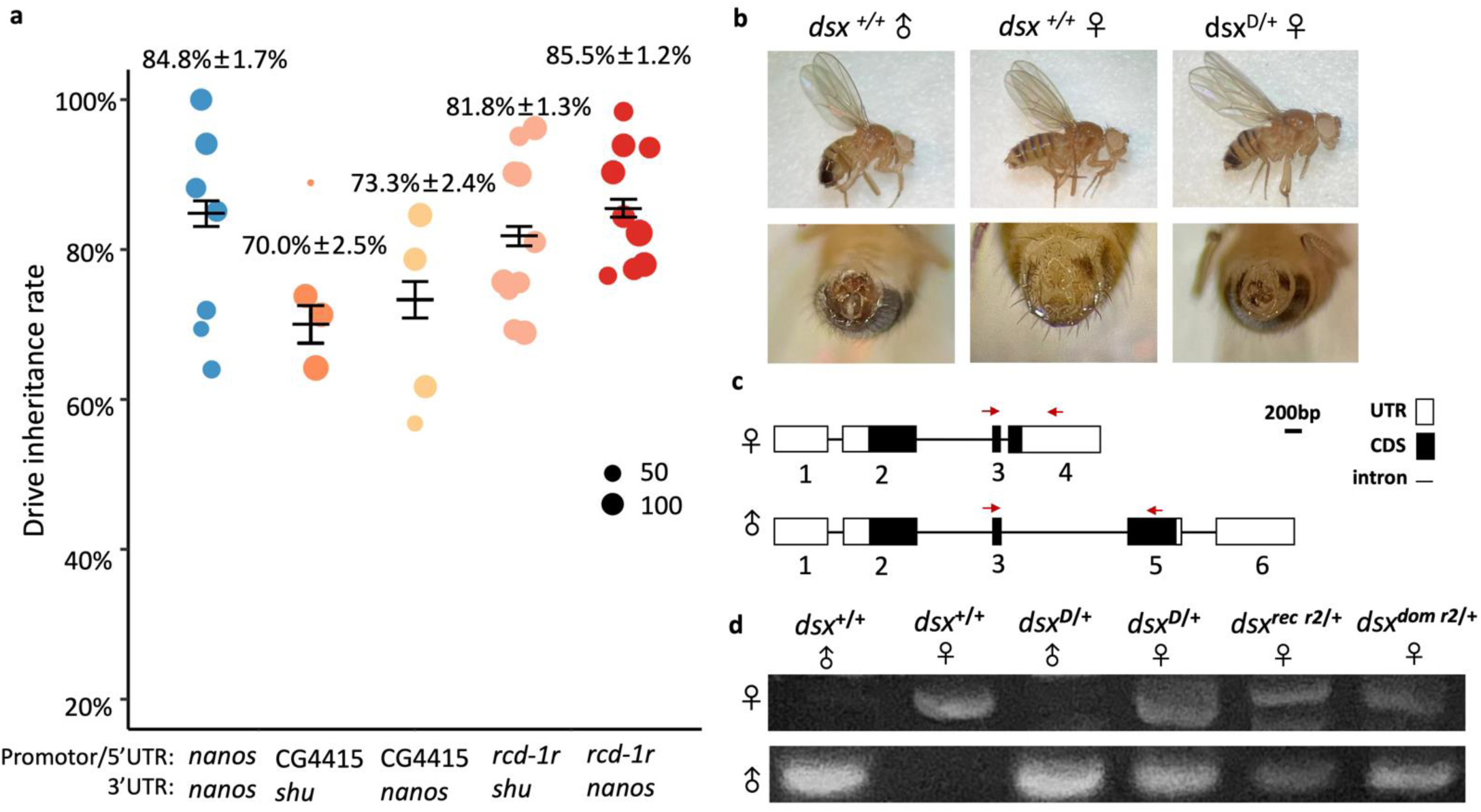
Drive inheritance, morphology of drive carriers, and *dsx* sex-specific transcript expression. (**a**) The drive inheritance rate of the progeny of *dsx* drive heterozygous males with one copy of Cas9 controlled by the indicated regulatory elements. Each dot represents progeny from a single drive male in one vial, and the size of each spot shows the number of individuals phenotyped. The mean inheritance rate (± s.e.m.) is shown. (**b**) Pictures of wild-type (dsx^+/+^) flies and female drive carriers (dsx^D/+^, and also carrying a *nanos*-Cas9 allele, though this does not change the phenotype) are shown. (**c**) Schematic representation of the male– and female-specific dsx transcripts. Red arrows are primers designed for male– or female-specific transcript diagnostic PCR. (**d**) Diagnostic PCR on cDNA using female-specific primers (top) and male-specific primers (bottom). dsx^rec r2/+^ refers to individuals with a recessive sterile r2 resistance allele, and dsx^dom r2/+^ refers to individuals with dominant sterile r2 alleles.

The DsRed in the eyes of drive individuals were observed to have two discrete levels of brightness. Bright DsRed individuals and faint individuals gave rise to progeny of the same brightness in the absence of Cas9. When Cas9 was present, faint individuals still had only faint offspring, but bright eye individuals had approximately one third of progeny that were faint eye, implying that these may have been the result of drive conversion. However, we found no sequence difference on drive construct and surrounding insertion site. For drive performance, we found that the faint eye males 75.7% drive inheritance with *nanos*-Cas9, representing a small but significant reduction (Fisher’s exact test, *P* < 0.0004, Supplementary Fig. 2). However, this apparently reduced drive inheritance may be partially due to the difficulty in phenotyping faint eye flies in the presence of EGFP.

Surprisingly, we found that all drive females were sterile, regardless of the presence of any Cas9 allele, indicating that the drive was dominant-sterile for females. Such females had a (partially) masculine phenotype, which is different from the *dsx*-target drive in *A. gambiae*, where the drive allele was recessive sterile, and heterozygous females lacked an abnormal phenotype, with only homozygous females showing a strong masculine phenotype (Kyrou et al. 2018). Examination of external sexually dimorphic structures in drive females showed phenotypic abnormalities, including a larger dark stripe at the end of the abdomen and male-like genitalia (Fig. 2b and Supplementary Fig. 1). To determine the molecular mechanism causing female-specific dominant sterility, we designed sex-specific primers and used cDNA as template for PCR (Fig.2c). We found that female– and male-specific versions of *dsx* transcripts were both expressed in drive females (Fig.2d). One possible reason is that the drive allele insertion disrupts the splice acceptor site of female-specific exon 4, preventing female-specific splicing from recognizing the site and recruiting splicing machinery. The next splice acceptor is then used instead, which is specific for the male version of the *dsx* transcript and results in incorrect sex development.

We also tested combinations of drives with different Cas9 lines varying in the amount of somatic expression^55^. Such somatic expression could give drive heterozygous males a similar phenotype to drive homozygous males, which could not be obtained because the drive is dominant female-sterile. Addition of Cas9 driven by the *nanos* promoter had no effect on the phenotype, as expected due to the very low somatic expression of *nanos*-Cas9. Compared to drive females carrying *nanos*-Cas9, drive females with Cas9 expressed by the CG4415 and *rcd-1r* promotors often had stronger masculine phenotypes (Supplementary Fig. 1, Supplementary Fig. 3).

### 3. Resistance allele formation and classification

To identify resistance alleles, which prevent Cas9 cleavage, we monitored the occurrence of mutations at the drive target site in the progeny of drive males with *nano*-Cas9. Though mutated *dsx* transcripts lose function, the *dsx* female transcript is haplosufficient. Yet strikingly, all resistance alleles had a dominant visible phenotype. In terms of morphological analysis and fertility, three types of nonfunctional (r2) resistance allele females were observed in our experiments: (1) dominant sterile r2 alleles with the same intersex/masculinized phenotype as drive females, which is the most common type of r2. (2) dominant sterile r2 alleles with a stronger intersex phenotype, representing about 8.3% of r2 alleles (Supplemental Data), and (3) recessive sterile r2 alleles with moderate intersex phenotype, but which were fertile when heterozygous for the r2 allele and a wild-type allele. The frequency of recessive sterile r2 alleles is about 5.3%, (2 of 38 in our r2 female fertility test, Supplemental Data). Females that were homozygous for this last class of r2 alleles displayed stronger intersex phenotypes. No functional/r1 resistance alleles were detected (wild-type progeny of drives males all had wild-type alleles), likely because the *dsx* target exon site was highly conserved and targeted by three gRNAs.

Diagnostic PCR suggested that male form *dsx* transcripts were generated both in dominant sterile as well as recessive sterile r2 females, but the male transcript expression level may be reduced in the latter (Fig. 2d). In all but the second class of resistance allele sequences, the region between the outermost gRNA cut sites were deleted. For r2 females with stronger intersex phenotypes, we saw a diverse mix of sequences, including full deletions between the outmost sites, individual indels that formed sequentially at each of the three gRNA target sites, and two sequences with an indel at the leftmost gRNA target site and incomplete homology directed repair of the drive after a simultaneous cleavage of the middle and rightmost gRNA target site (Fig. 3). Remarkably, the AG in the 3’ intron, the splice acceptor site, was preserved in all but one sequence for females with moderate intersex/masculinized phenotype, while AG was partially or fully deleted in females with stronger intersex phenotypes. Thus, it is possible that the disruption of the splicing acceptor site in female-specific exon 4 increased the possibility of generating more male-specific *dsx* products. In contrast, the preservation of the splice acceptor site enabled (disrupted) female *dsx* products to be generated in spite of mutations ^52^, reducing but not eliminating the chance that splicing skipped directly to male-specific exon.

**Figure 3.**
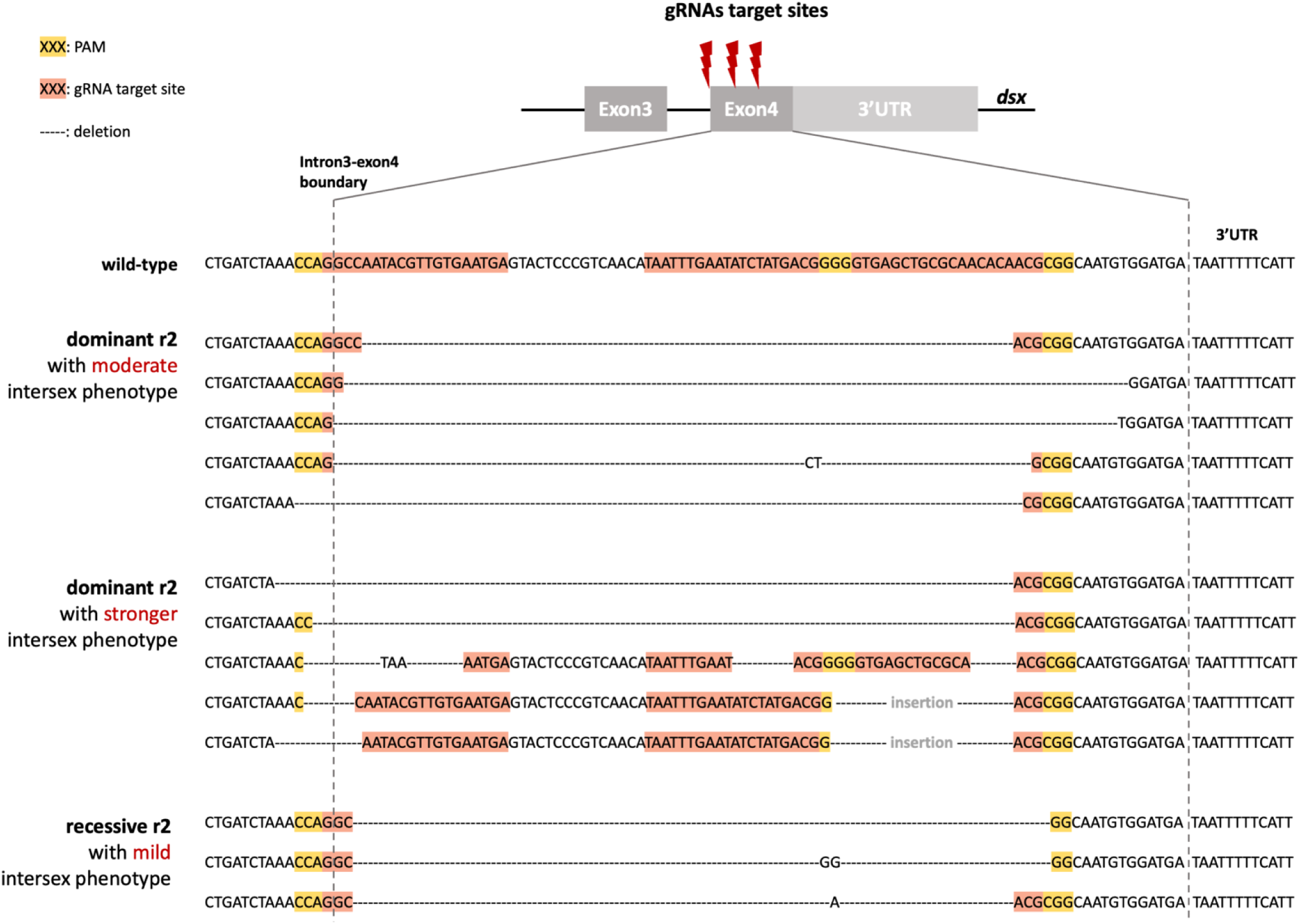
Sequences of three types of nonfunctional resistance alleles. Each sequence originates from one female carrying a resistance allele with the listed phenotype. Vertical dashed lines indicate the boundaries of the female-specific exon’s coding sequence. Orange highlighting show the gRNA target sequences, and yellow show the gRNA PAM sequences. ––– indicates deletion. “Insertion” refers to a 311 bp region from the right side of the drive that was copied by incomplete homology-directed repair.

### 4. New pest control strategy RIDD system

Based on the dominant-sterile property of *dsx* drive in *Drosophila*, the largely dominant-sterile resistance alleles, and the high total germline cut rates in males, we put forward a new pest suppression strategy called the Release of Insects carrying a Dominant sterile Drive allele (RIDD) system. Drive conversion can take place in germline of male heterozygotes, which are released continuously in large number into a population, similarly to the RIDL strategy. When drive males are released to wild population, the drive allele can be transmitted when they mate with wild-type female. Nearly all female progeny will be sterile (regardless of whether they inherit drive alleles or resistance alleles), yet will still contribute to resource competition. The drive male offspring will usually be able to continue “homing”, but if they don’t receive a drive, they can still pass on a dominant-sterile resistance allele to half of their progeny (Fig. 4a). Repeatedly releasing drive males into a wild population will result in a steady increase in the drive allele frequency. Eventually a sufficient number of females will be sterile, and the wild population will be suppressed (Fig. 4b). This system combines the merits of gene drive and RIDL. It is more powerful and efficient than fsRIDL, but still self-limiting, allowing for highly confined population suppression compared to standard homing suppression drive systems.

**Figure 4.**
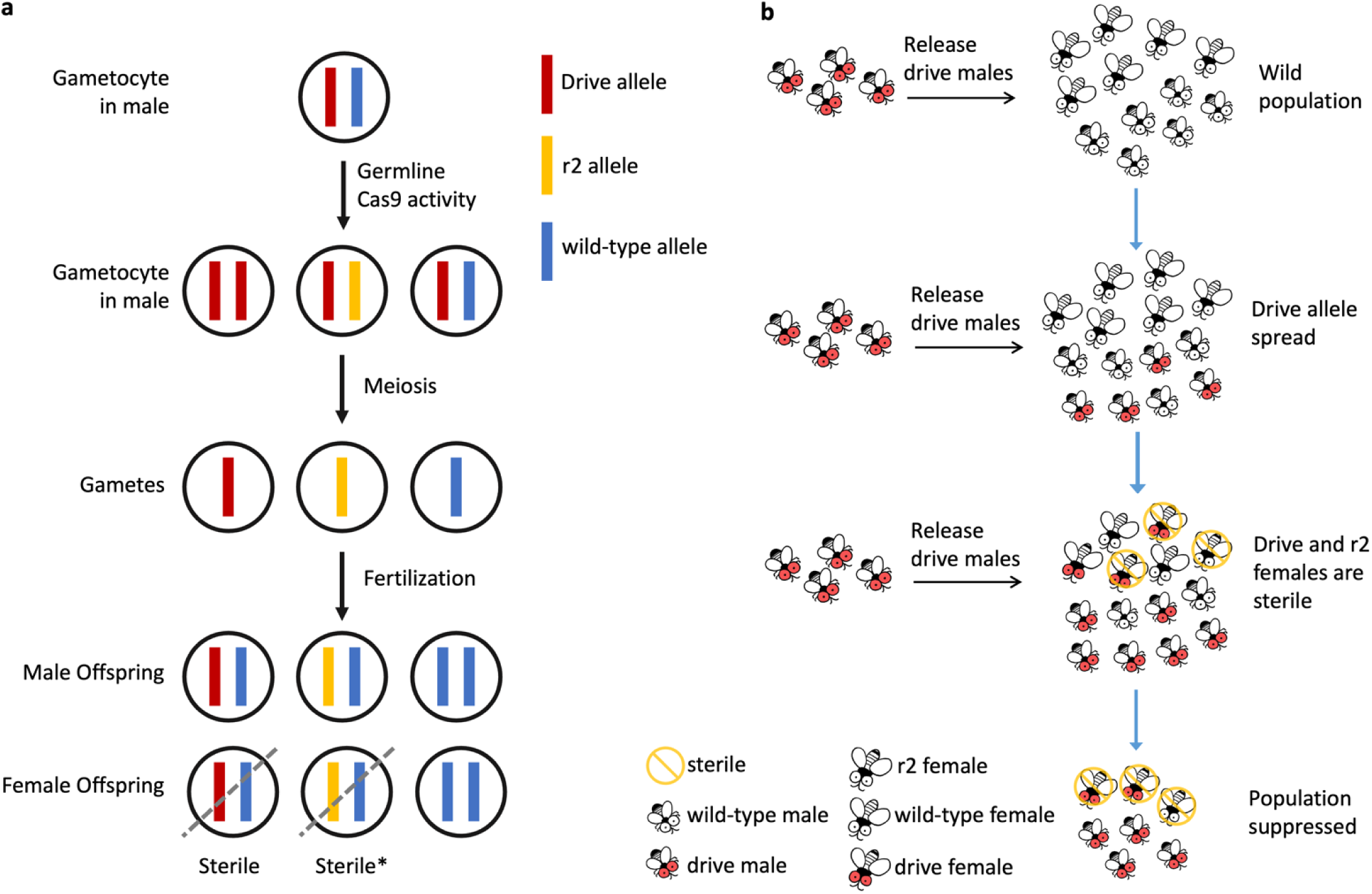
Scheme of Release of Insects carrying a Dominant sterile Drive (RIDD) system. (**a**) Drive inheritance of RIDD. In male drive heterozygotes, germline Cas9 activity converts wild-type allele to drive alleles by homology-directed repair, while end joining repair or incomplete homology-directed repair generates nonfunctional resistance (r2) alleles. Females carrying one drive allele are sterile. The most common type of r2 is also dominant sterile in female, though a small fraction are recessive sterile. A small fraction of wild-type alleles in the male germline may also remain uncut. (**b**) Concept of RIDD pest control strategy. Drive heterozygous males are continuously released into a wild population. Drive conversion and resistance allele formation take place in the male germline, so nearly all female progeny of drive males are sterile (drive carrier and most r2 carrier females are sterile). Over time, the frequency of drive carriers increases, and with a high enough release level, the population will eventually be suppressed.

### 5. Assessment of the RIDD system in cage populations

To test the Release of Insects carrying Dominant sterile Drive (RIDD) system, we carried out a cage study. We first conducted male mating competitiveness assay, so as to test the fitness of drive males compared to wild-type males, a critical performance parameter for systems involving continuous male releases. In 1:1 competitions, drive males successfully mated in 39 of 74 tests, while wild-type males successfully mated in 31 tests. In four vials, both males mated with the female, based on the phenotype of the offspring. There was no significant difference between the mating competitiveness of drive males and wild type males. Then, two replicate cage studies were carried out. Eight weeks were taken to establish an initial population for the cages, which was approximately 2500. Overlapping-generation cages were maintained by adding fresh food every few days and removing old food. The population size was measured each week (Fig. 5a). To assess the spread of the drive, flies in cages were randomly aspirated for phenotyping (Fig. 5b and Supplementary Fig 4.a) and then put back into the cage. In addition, to monitor the ratio of drive carriers among newly hatched progeny, the old food bottles were often kept for extra one day outside cages, and then newly emerged adulted were phenotyped (Supplementary Fig. 4b).

**Figure 5.**
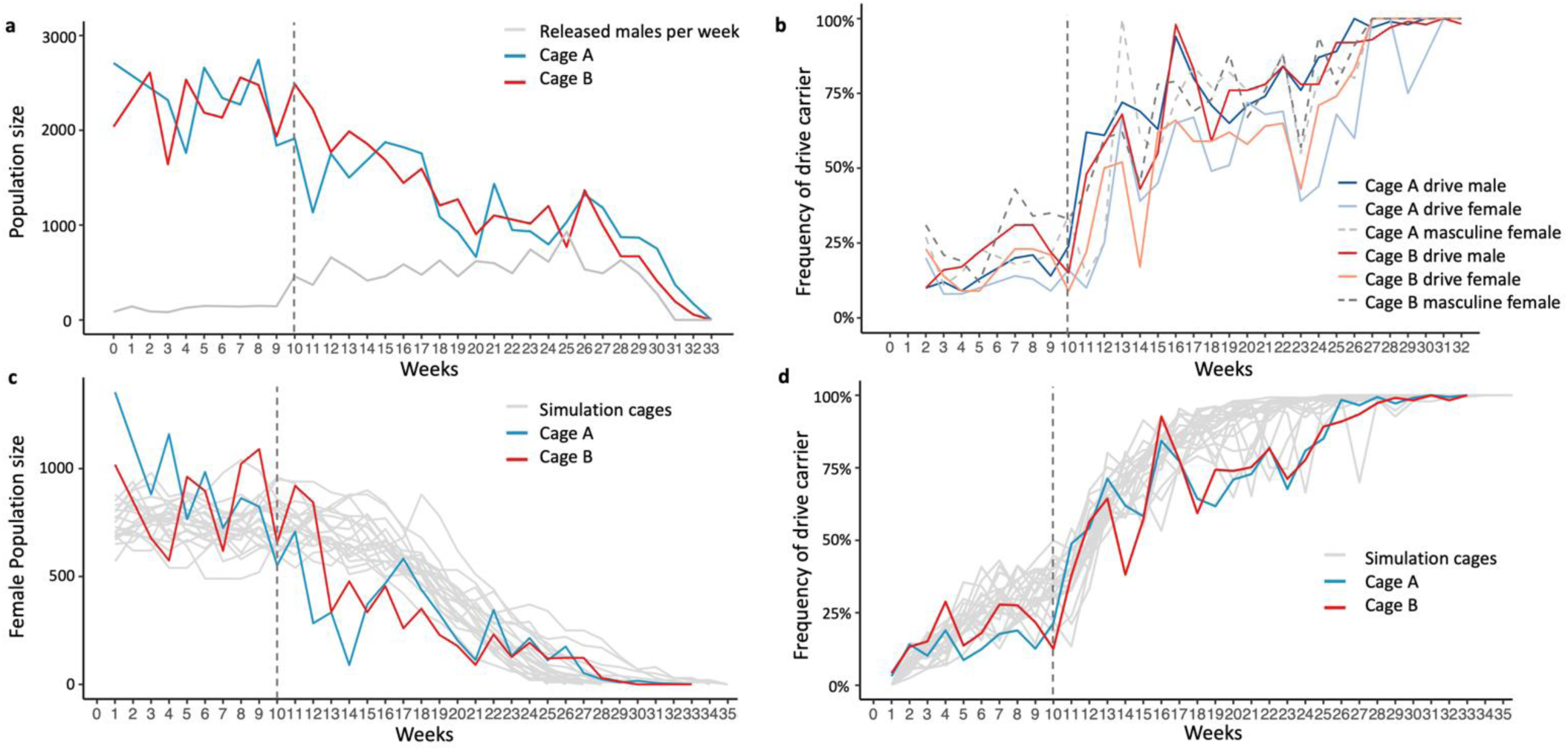
**Population dynamics of cages**. *dsx*^D/+^ males were released each week (usually at two evenly spaced time point per week) into continuously maintained fly populations with overlapping generations. (**a**) Population size of cages A and B together with the number of males released in each week. After the weekly release size was increased, the population steadily declined. (**b**) Frequency of individual phenotypes the cages based on a small weekly sample of flies. Masculine phenotype females include both sterile drive females and those with a nonfunctional resistance allele. (**c**) Female population size in cages. This data was in agreement with stochastic simulations of the cage population (grey lines). (**d**) Frequency of drive individuals in cages A and B together with results from twenty simulations. The vertical dashed line marked the week where the drive male release size was increased.

In the first ten weeks, we released 82∼148 *dsx* drive males into each cage each week, which was about 3∼7% of total population (Fig. 5a). We did not observe a significant decrease in the population size, and the frequency of drive carriers appeared to stabilize at a low level (Fig. 5a). However, the existence of drive females verified that released males mated successfully in the cage population. Starting from week 10, the release size of *dsx* males was substantially increased to approximately 500 per cage each week, which was about 20% of total population at first (Fig. 5a). We observed gradually increasing ratios of drive carriers in cages and among newly hatched individuals (Fig. 5b, Supplementary Fig. 4b). In the later period of the study, the drive carrier frequency reached 100% in both cages. We observed the sex ratio biased toward males, and the population size steadily decreased in both cages (Fig. 5a, Supplementary Fig. 4b). The release size in the later period of cage study was ∼50% of the total population. No new hatched individuals were produced in both Cage A and B, and the cage collapsed at week 29. At this time, we stopped releasing drive males. At week 32, we collected all the remaining flies for phenotyping and stopped the cages. All flies in cage B were drive males, while all flies in cage A were drive males except two r2 females with strong intersex phenotype.

For comparison, we also maintained a control cage for eleven weeks (Supplementary Fig. 5) with an initial 300 Cas9 flies. The population size gradually increased and achieved an equilibrium at about week 4, which was approximately 1200 flies.

SLiM was used to simulate the population dynamics of our RIDD cage experiments, using estimates for several parameters related to the species-specific ecology of the laboratory cage populations. We assumed the same drive conversion rate in our experiments, no fitness costs, and no remaining wild-type alleles in the germline of male drive heterozygotes, with all resistance alleles being the dominant-sterile type. The trajectories of female population size and drive carrier frequency of twenty simulations are closely matched to the experimental cage study, indicating that the system performed as expected with no major undetected fitness costs.

### 6. Modelling of RIDD system

For RIDD and similar technologies such as SIT and RIDL, success depends on not just the drive performance and release parameters, but also heavily on the ecology of the species, in particular the density growth curve. The low-density growth rate (relative growth in the absence of competition) is important, but the shape of the curve is also essential. This is because repeated releases will give an increased release ratio of transgenic to wild-type males as the wild-type population shrinks. Thus, the release size does not need to overcome the low-density growth rate when the wild-type population is at equilibrium, only reduce it enough for the male transgenic ratio to eventually reach this level. However, this complexity can make it difficult to compare different transgenic constructs in the general case.

Here, we introduce a possible method of comparison based on genetic load (the fraction of females that don’t successfully reproduce, in this case defined as having a normal number of fertile daughters) to assess the suppressive power of the RIDD system. We developed a deterministic, discrete-generation model framework. We assume that the wild-type population does not change, and thus, the ratio of released males to native-born males remains constant. We then assess the genotype of the population for several generations until equilibrium is reached. At this point, we report the genetic load. Because the total cut rate was high in our experiments and almost all resistance alleles were dominant-sterile, we assume that any wild-type allele in the germline of a male drive heterozygote will be converted to a dominant-sterile resistance allele if it is not converted to a drive allele. By varying the drive conversion and release ratio, we see that higher drive conversion notably improves the performance of the system (Fig. 6a). However, even with no drive conversion, the system is quite efficient compared to RIDL (where released males are homozygous) and especially classic SIT.

**Figure 6.**
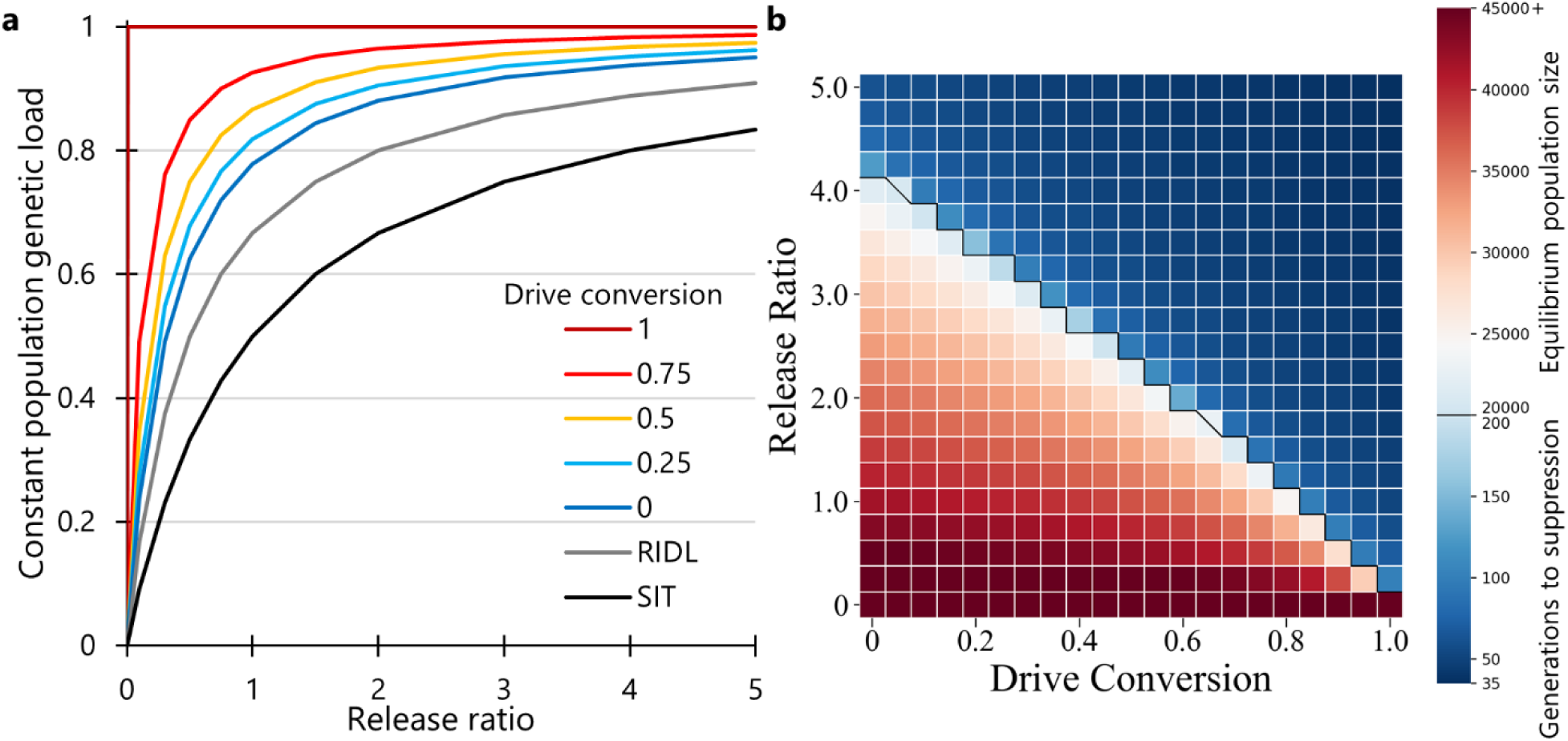
**Modeling the performance of the RIDD system**. A constant number of drive heterozygous males containing a RIDD system were released each generation at with the number based on the specified ratio of released males to males in the starting population. (**a**) The genetic load at equilibrium (after several generations of drive releases results in a constant drive frequency) for varying drive release ratio and drive conversion rate. The model is deterministic and uses discrete generations All wild-type alleles in the germline not converted to drive alleles are assumed to be converted to dominant-sterile resistance alleles. SIT and RIDL are shown for comparison. RIDL releases are homozygous males. (**b**) The heatmap shows the outcome of stochastic simulations for suppression of a population of 50,000 with discrete generations and a linear density growth curve. The lower end of the heat bar shows the average number of generations to suppression, and the upper portion shows the average native population size after 200 generations, representing the equilibrium population size. The black line shows the boundary for these two outcomes, but for two spots in the parameter space (with diagonal lines through them), population elimination occurred in an intermediate fraction of the simulations (above 0, but below 1). Each point in the parameter space had 20 replicates.

To model the success of RIDD in large, panmictic populations, we used our SLiM framework to simulate populations averaging 50,000 individuals with a low-density growth rate of 10 and a linear density-dependent growth curve. This creates a highly robust population, substantially more so than our cage population. In this case, high drive conversion was very useful for successful population suppression, proportionally reducing the required release size (Fig. 6b). When there were over four males released for each male in the native starting population, the system could be successful even without drive conversion.

## Discussion

In this study, we proposed a new pest control strategy, Release of Insects carrying a Dominant sterile homing gene Drive (RIDD). This system was tested by targeting *doublesex* in *D. melanogaster*. In our drive design, three gRNA were used to target the 5’ intron-exon boundary of the female-specific exon, a highly conserved region. The drive has high inheritance bias in heterozygous drive males, which were healthy and fertile, like a standard homing gene drive. However, all female drive heterozygotes and nearly all female resistance allele heterozygotes were sterile with a masculinized phenotype. This result is in contrast to recent studies in *A. gambiae* and *D. suzukii* that also target *doublesex* ^40, 56^. Their systems are also homing suppression gene drives, but females are sterile only when they are homozygous with drive alleles in the *A. gambiae* drive. In the *D. suzukii* study, some drive alleles tested are dominant female-sterile, but not the resistance alleles like in our study. Our sex-specific transcripts analysis suggests that dominant sterility can be accounted for incorrect splicing in females. Male and female *dsx* transcripts are co-expressed in drive and r2 females, implying that drive allele insertion breaks the splice acceptor site of female-specific exon 4 and prevents female-specific splicing components from recognizing the site. However, instead of just skipping the splicing and disrupting the female version, splicing directly skips to next splice acceptor in front of male-specific exon and therefore generates the male version of the *dsx* transcript. This leads to a sex development disorder in females. In the *dsx*-target gene drive study in *D. suzukii*, the site they chose to cut was fully within the female-specific exon, preserving the splicing acceptor site, so the transcript of the disrupted allele is allowed to splice correctly in spite of at least some end-joining mutations^56^. Though one allele loses its function, the other copy of functional *dsx* is sufficient for female sex development in the absence of the male splice form. However, this explanation is not completely clear on why drive alleles from this site are dominant-sterile. Perhaps, nucleotides in the exon can be required for splicing even toward the middle of the exon, not just those immediately adjacent to the intron-exon boundary. The level of skipping to the male splice form is would be a matter of degree, with higher levels of splicing disruption with increasing sequence disruption throughout the first half or more of the exon. This is suggested by our detailed analysis of nonfunctional resistance alleles.

Three types of non-functional resistance (r2) alleles were detected on the basis of masculine phenotype intensity and fertility: dominant sterile r2 females with stronger masculine phenotype, dominant sterile r2 females with weaker phenotype (the same phenotype as drive alleles in the absence of somatic Cas9 expression), and recessive sterile r2 females with the same weaker phenotype. We further found that this different degree of masculinization is closely related to the degree of preservation of the AG in the intron-exon boundary (the splice acceptor site). In r2 alleles preserving AG after end-joining, less male *dsx* products will be generated, resulting in a milder phenotype (even more nucleotides being preserved potentially allow for a recessive sterile phenotype, in addition to being visually milder). All types of r2 alleles are soon removed from the population in sterile females, and this process is much faster for the more common dominant-sterile alleles. This result promises a strong population suppression power for homing drives as its frequency reaches equilibrium, because most remaining females are sterile. Dominant-sterile r2 alleles will not slow population suppression, which is a problem encountered by recessive-sterile r2 alleles^57^.

Our overlapping-generation cage study showcased the feasibility of RIDD in population suppression as well as demonstrating a relatively easy model by which cage studies with overlapping generations could be conducted (which is potentially more representative of application scenarios than more commonly used discrete generations). With weekly releases of drive males increasing to approximately 50∼60% of the equilibrium male population size, the population was successfully eliminated in two replicate cages. The RIDD system is similar to combining fsRIDL with gene drive, which was originally shown to substantially increase RIDL’s efficiency ^20^. Specifically, homing takes place in drive male germline, biasing the allele’s inheritance, which maintains the RIDL allele in the population for a longer time. Compared to fsRIDL, RIDD is substantially more efficient and powerful, as indicated by our modeling results. Even with homing, RIDD has the advantage of sterilizing females with r2 alleles, not just with drive alleles. However, maintaining the drive is more complex than RIDL systems. Though there is no need for an antibiotic repressor, drive males must continually be crossed to wild-type females to keep the drive line.

Nevertheless, though more powerful than similar alternatives, the RIDD system is still self-limiting. If releases are ceased at any time before the population collapse, the population will easily recover because drive females produce no progeny, so the drive frequency will decline and be eliminated in several generations. This property allows for highly confined population suppression compared to standard homing suppression drives.

Additionally, apart from the *nanos* promoter for Cas9, the *CG4415* promotor, *rcd-1r* promotor, and *shu* 3’UTR were tested for Cas9 constructs in our study. Cas9 guided by *rcd-1r* has similar drive conversion as Cas9 regulated by *nanos*. Both *CG4415* and *rcd-1r* promotors contribute to higher somatic expression, reflected in a higher proportion of stronger masculine phenotypes among drive carrier offspring (Supplemental Data, Supplementary Fig.4). For RIDD, the property of somatic expression is acceptable because it will not affect males, and drive females are sterile anyway. Indeed, a standard homing suppression drive that does not have any dominant-sterile r2 alleles could still be used for similar suppression if somatic Cas9 expression rendered drive alleles dominant sterile, though such a drive would be less efficient than RIDD due to weaker contribution to population suppression from r2 alleles.

Overall, our study suggests some mechanisms and possible approaches for dominant sterility and drive target site design for *dsx*. Because the *doublesex*-based sex determination pathway is highly conserved in dipterans, including mosquitos, moths, and flies, our design utilizing *dsx* sex-specific splicing with dominant effects has the potential to be applied to a wide variety of pest species control when splicing follows the paradigm in *Drosophila* rather than *Anopheles*. Moreover, the RIDD pest control system, verified by our modeling and cage experiments, combines the merits of high efficiency and self-limiting, which will be a promising pest control strategy for practical application.

## Acknowledgements

Thanks to Jie Du, Jingheng Chen, Xuqing Feng, and Weiwen Yang for assistance with cage experiments. This study was supported by laboratory startup funds from Peking University, the Center for Life Sciences, and the NSFC Overseas Youth Fund.

## SUPPLEMENTARY INFORMATION

### Supplementary methods

#### Plasmid Construction

Plasmid with dsx gRNAs: HSDdsx3g

Plasmid with dsx Left-right arm: HSDdsxHA

Final donor plasmid: HSDdsxRed3g

|  | *Template* | *Primer/Enzyme 1* | *Primer/Enzyme 2* |
| --- | --- | --- | --- |
| **HSDdsx3g** |  |  |  |
| *PCR product* | TTTgRNA<sup>tRNAi</sup> | dsx_g31_F | dsx_g31_R |
| *PCR product* | TTTgRNA<sup>t</sup> | dsx_g12_F | dsx_g12_R |
| *PCR product* | TTTgRNA<sup>t</sup> | dsx_g23_F | dsx_g23_R |
| **HSDdsxHA** |  |  |  |
| *PCR product* | w<sup>1118</sup> Genome | dsxLeft_F | dsxLeft_R |
| *PCR product* | w<sup>1118</sup> Genome | dsxRight_F | dsxRight_R |
| *PCR product* | HSDygU4 | backbone_F | backbone_R |
| **HSDdsxRed3g** |  |  |  |
| *Plasmid Digest* | HSDdsxHA | EcoR1 | Xmal |
| *PCR product* | HSDdsx3g | dsx3gRNAs_F | dsx3gRNAs_R |
| *PCR product* | HSDygU4 | dsx_dsRed_F | dsx_dsRed_R |

#### Construction primers

**HSDdsx3g**

dsx_g31_F: GTGCATAATTTGAATATCTATGACGGTTTTAGAGCTAGAAATAGCAAGTTAAA

dsx_g31_R: AAAACGGCCAATACGTTGTGAATGATGCATCGGCCGGGAATCG

dsx_g12_F: GCATCATTCACAACGTATTGGCCGTTTTAGAGCTAGAAATAGCAAGTTAAA

dsx_g12_R: AACCGTTGTGTTGCGCAGCTCACTGCACCAGCCGGGAATCG

dsx_g23_F: GCAGTGAGCTGCGCAACACAACGGTTTTAGAGCTAGAAATAGCAAGTTAAA

dsx_g23_R: CGTCATAGATATTCAAATTATGCACCAGCCGGGAATCG

**HSDdsxHA**

dsxLeft_F: AATAGGGGTCAGTGTGGAATGTATCTCATGTGGCCCA

dsxLeft_R: GCGCAGCTCCCGGGGGGGAATTCGCCTGGTTTAGATCAGATCGAGAG

dsxRight_F: ACCAGGCGAATTCCCCCCCGGGAGCTGCGCAACACAACGC

dsxRight_R: ACTAAGCAGAAGGCCATCTCAGCGAGGCCGATTTT

backbone_F: CGGCCTCGCTGAGATGGCCTTCTGCTTAGTTTGATG

backbone_R: CATGAGATACATTCCACACTGACCCCTATTTGTTTATTTT

**HSDdsxRed3g**

dsx3gRNAs_F: CATCAATGTATCTTA TTTTTTGCTCACCTGTGATTGCT

dsx3gRNAs_R: ATCCACATTGCCGCGTTGTGTTGCGCAGCTCATGCATACGCATTAAGCGAACA

dsx_dsRed_F: GAATGTCTCTCGATCTGATCTAAACCAGGCGGCGGCCGCGGATCTAATT

dsx_dsRed_R: CAGGTGAGCAAAAAATAAGATACATTGATGAGTTTGGACA

#### Diagnostic PCR for dsx transcripts primers

**Female specific primer pairs**:

dsx_exon3_S_F: TTGGGCCAAGACGTTTTCCT

dsx_exon4_S_R: CGTTGATTTGGCGTGCGTAT

Expected Length: 713bp

**Male specific primer pairs:**

dsx_exon3_S_F: TTGGGCCAAGACGTTTTCCT dsx_exon5_S_R: AGGGCTTTTGGTTGTGGACA

Expected Length: 382bp

## Supplementary Figures

**Supplementary Figure 1.**
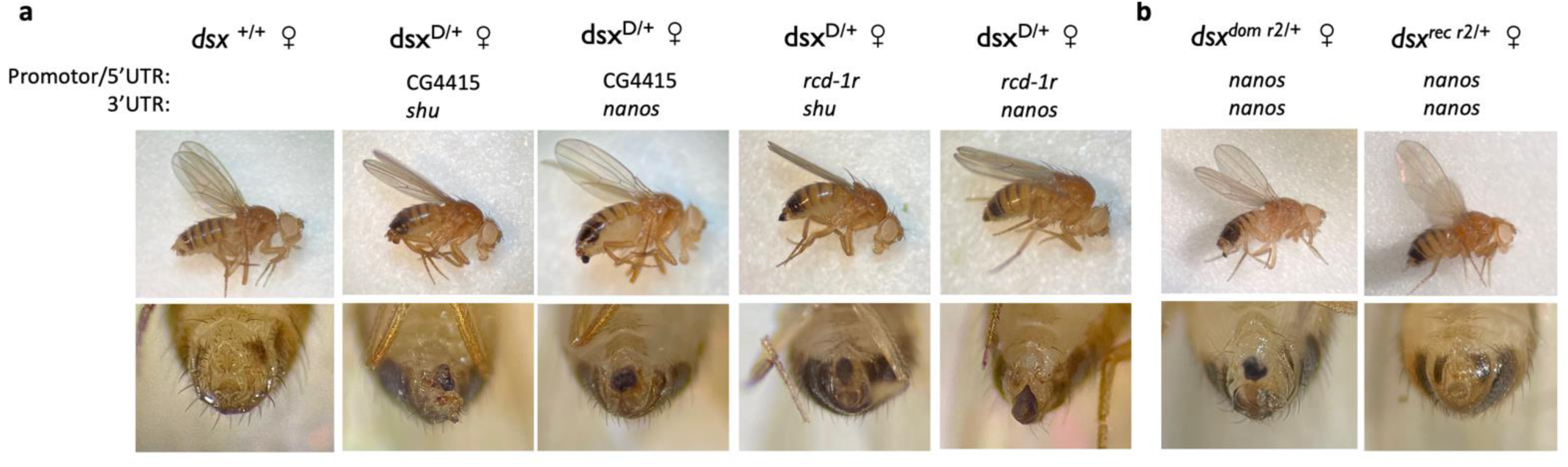
Additional pictures showing female morphological phenotypes. **(a)** Drive females (dsxD/+) carrying Cas9 with different promotor/5’UTR and 3’UTR combinations. Wild-type (dsx^+/+^) are included as a control for comparison. Note the stronger phenotype in females with the *CG4415* and *rcd-1r* Cas9 promoters (these each have moderate somatic expression, while *nanos* had little to none). (**b**) Two types of nonfunctional resistance allele females. dsx^dom r2/+^ refers to dominant sterile r2 individuals, dsx^rec r2/+^ refers to recessive sterile r2 individuals. Both of dsx^dom r2/+^ and dsx^rec r2/+^ were homozygous for *nanos*-Cas9-*nanos* allele.

**Supplementary Figure 2.**
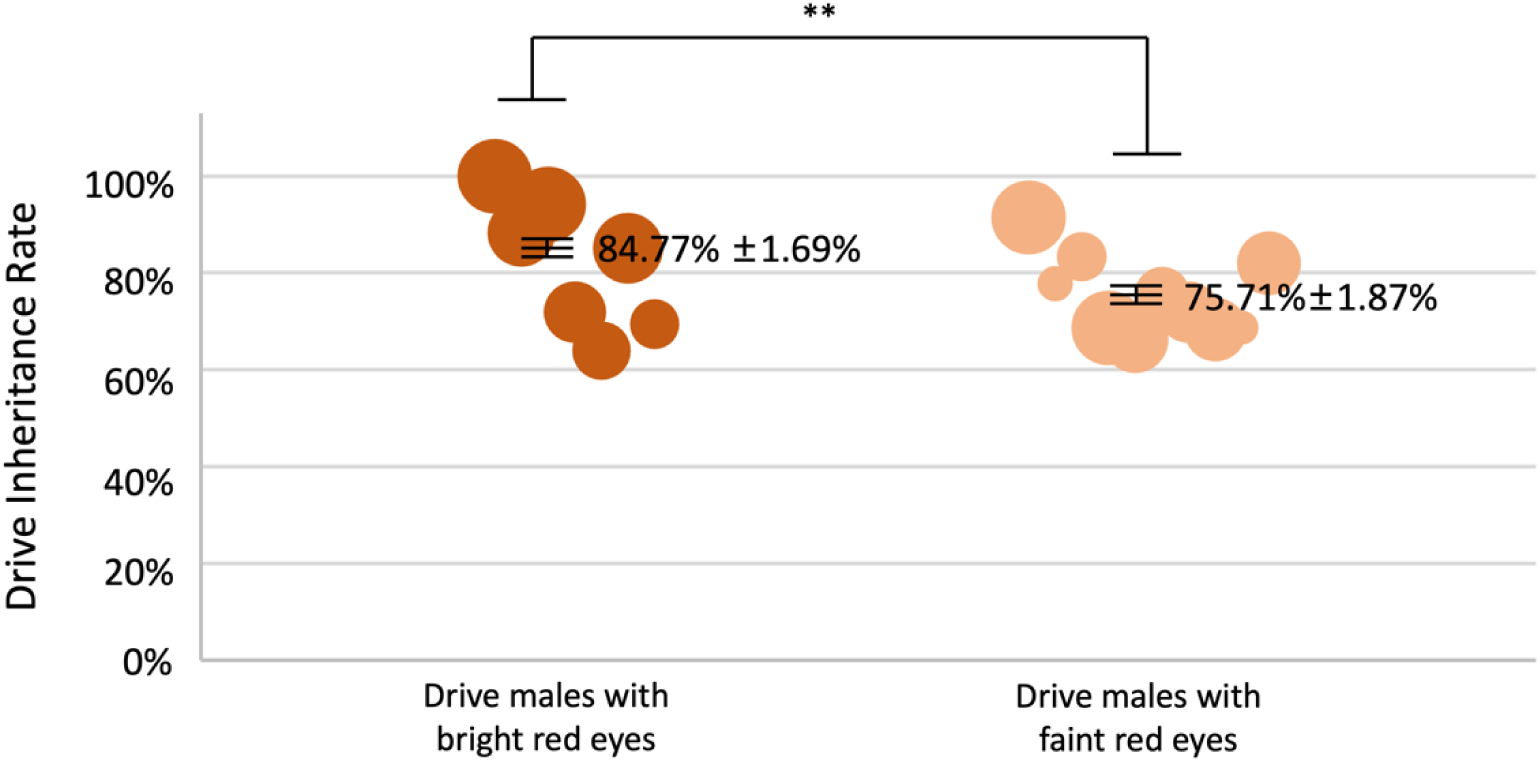
Drive inheritance rate of drive males with bright eyes and faint eyes. The drive inheritance rate of the progeny of *dsx* drive heterozygous males that were homozygous for *nanos* Cas9. Each dot represents progeny from a single drive male in one vial, and the size of each spot shows the number of individuals phenotyped. The mean inheritance rate (± s.e.m.) is shown.**\*\*** represents P<0.0004 in Fisher’s exact test.

**Supplementary Figure 3.**
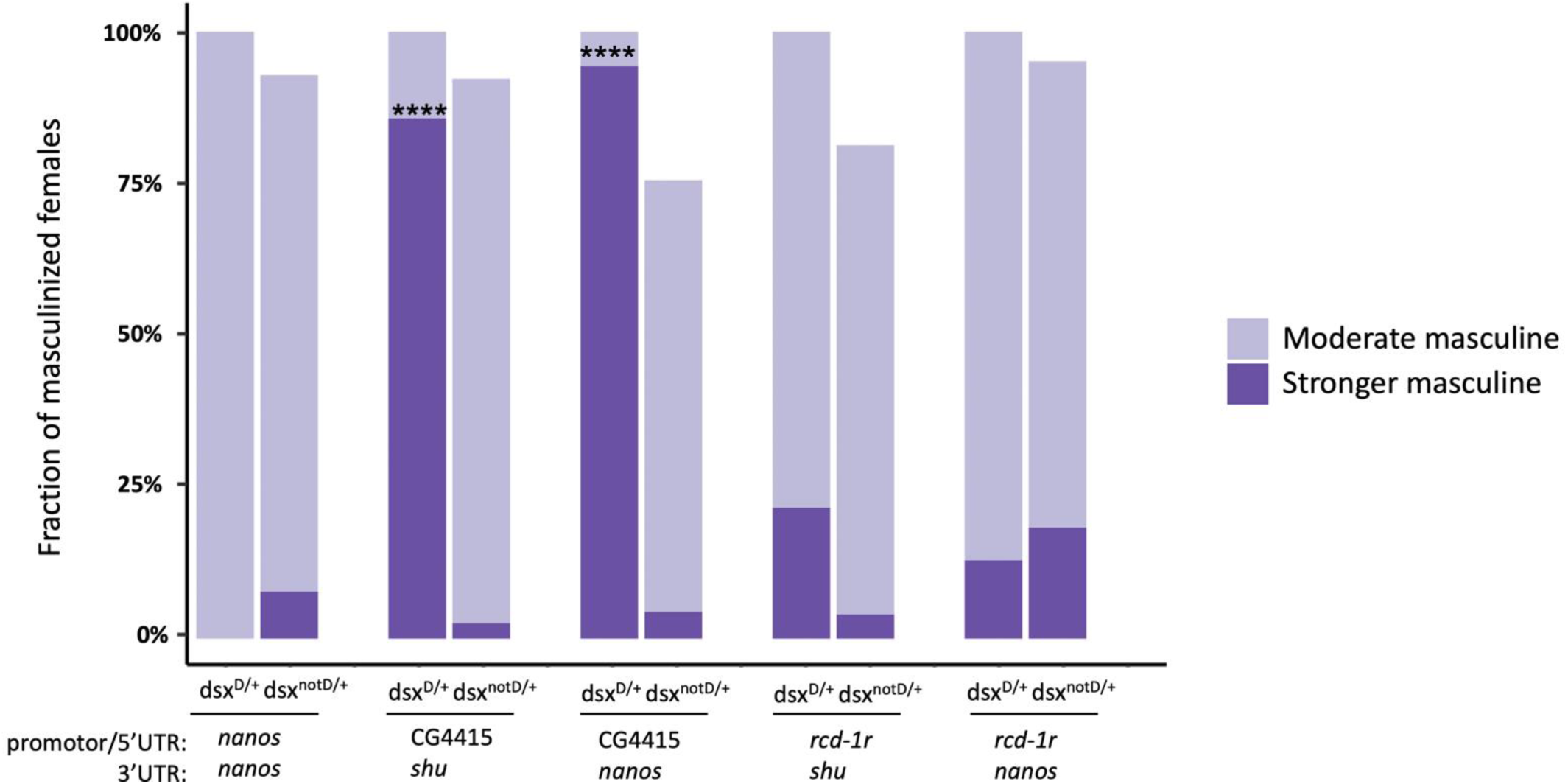
Somatic expression of *dsx* drive females with different Cas9 cassette. Female Cas9-bearing progeny of males with the drive and Cas9 were phenotyped. The bars show the fraction of females with the indicated phenotype out of all those with the drive (left bar) or without the drive (right bar). The stronger phenotype is most likely a result of somatic Cas9 expression. Non-drive females that lack any masculine phenotype most likely received an intact wild-type allele. Lack of a drive allele prevents cleavage in the soma, so the stronger phenotype was not observed in these females (all their cells would always have an uncut wild-type allele).

**Supplementary Figure 4.**
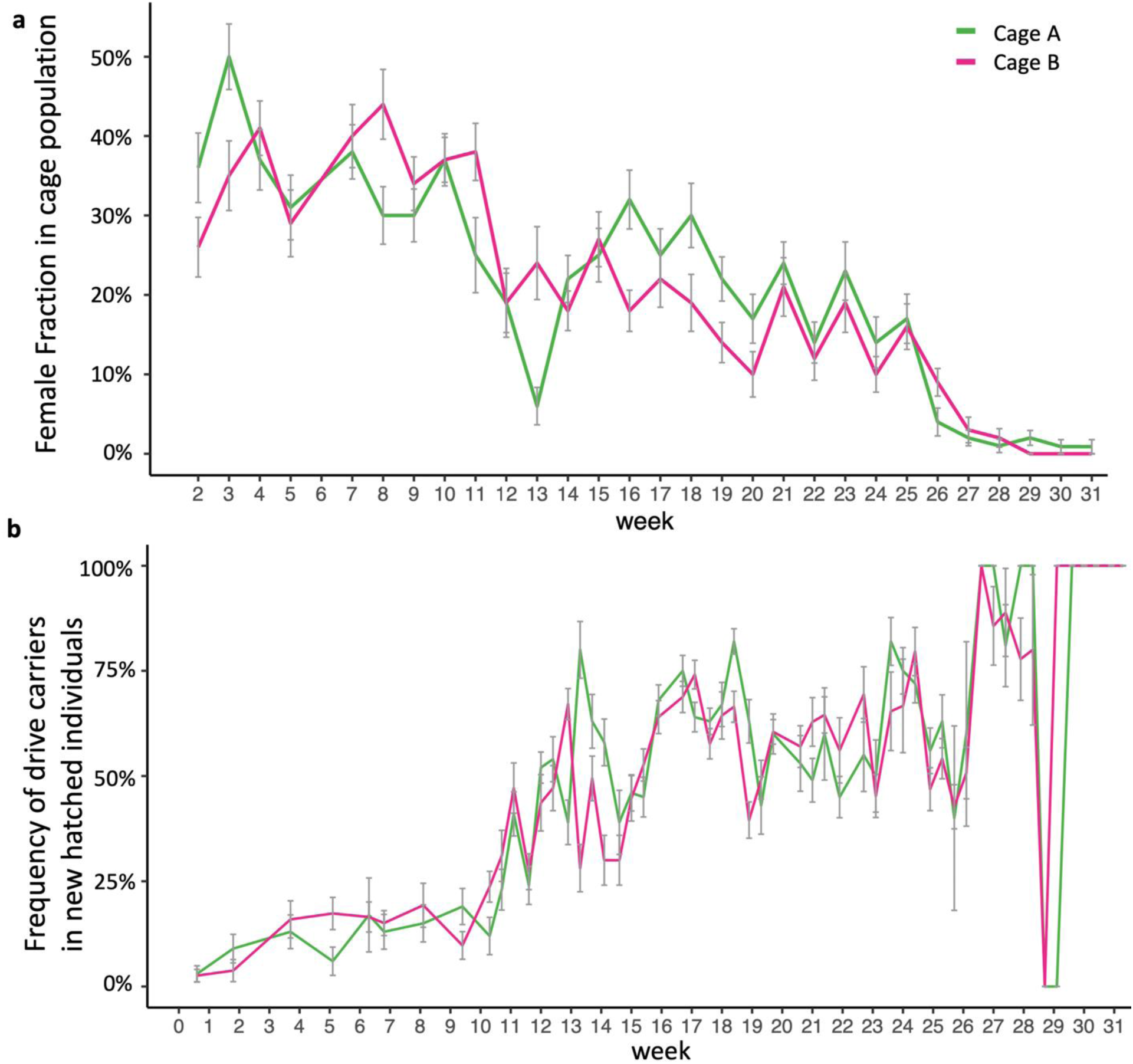
Sex Ratio and progeny drive carrier frequency in cage populations. (**a**) The fraction of females in the cages. The cages showed a male bias due to increased female mortality during egg-laying and continuous release of drive males. (**b**) Frequency of drive carriers that hatched from cage bottles one day after the bottle was removed from the cage. Grey bars indicate the standard error of the mean.

**Supplementary Figure 5.**
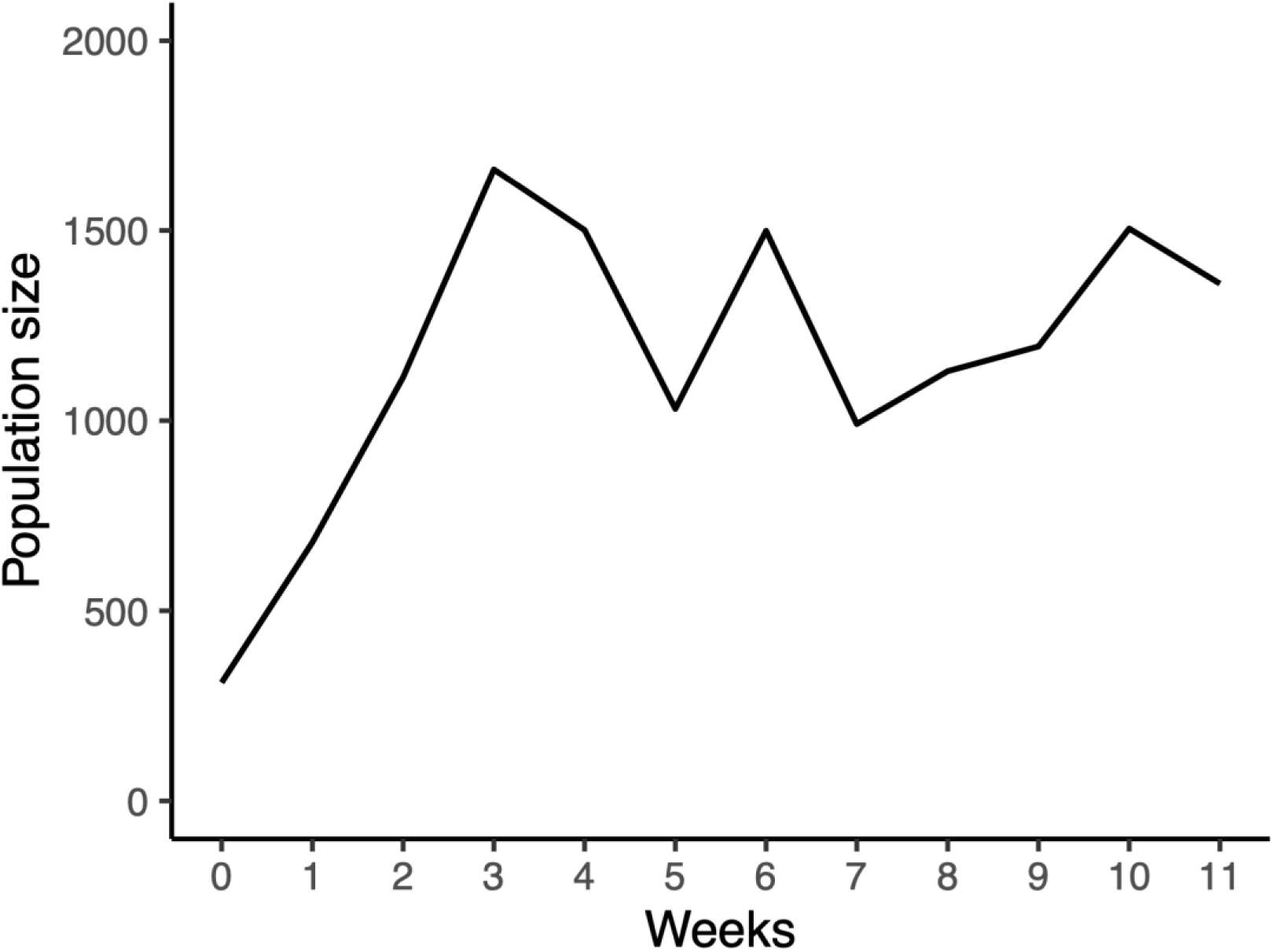
Population size of overlapping-generation control cage. A control cage with an initial 300 *nanos*-Cas9 homozygous flies and 200 flies released two week later was maintained for 11 weeks. The cage was treated in the same way as the experimental cages with five food bottles in circulation, replacing one every three days. The equilibrium population size was approximately 1200 flies for this time period, which was reached after about three weeks.

## Notes

https://github.com/jchamper/ChamperLab/tree/main/dsx-Suppression-Drive

### Competing Interest Statement

The authors have declared no competing interest.

